# Comprehensive analysis of network reconstruction approaches based on correlation in metagenomic data

**DOI:** 10.1101/2023.06.20.545665

**Authors:** Alessandro Fuschi, Alessandra Merlotti, Thi Dong Binh Tran, Hoan Nguyen, George M. Weinstock, Daniel Remondini

## Abstract

Microbiome analysis is transforming our understanding of biological processes related to human health, epidemiology (antimicrobial resistance, horizontal gene transfer) environmental and agricultural studies. At the core of microbiome analysis is the description of microbial communities based on quantification of microbial taxa and dynamics. In the study of bacterial abundances, it is becoming more relevant to consider their relationship, to embed these data in the framework of network theory, allowing characterization of features like node relevance, pathway and community structure. In this work we characterize the principal biases in reconstructing networks from correlation measures, associated with the compositional character of relative abundance data, the diversity of abundances and the presence of unobserved species within a single sample, that might lead to wrong correlation estimates. We show how most of these problems can be overcome by applying typical transformations for compositional data, that allow the application of simple measures such as Pearson’s correlation to correctly identify the positive and negative relationships between relative abundances, when data dimensionality is sufficiently high. Some issues remain, like the role of data sparsity, that if not properly addressed can lead to imbalances in correlation coefficient distribution.

## Introduction

Techniques based on next-generation sequencing (NGS) can elucidate the complex functioning of natural microbial communities directly in their natural environment. New branches of research have been created such as the study of the human microbiota which showed heterogeneity between different anatomical sites and individual variablity [1, 2], or the ability to characterize and monitor the presence of AMR (antimicrobial resistance) worldwide [3]. Together with the analysis on sample microbiome composition, it is also useful to investigate the relationship between the observed species or OTUs (Operational Taxonomic Units, which from now on will be defined as taxa). Network theory provides many essential tools to characterize macro properties of the ecology of a natural environment by defining central elements or communities in the system and allowing visualization of these results by exploiting network structural properties [4]. Therefore, the first step to reconstruct any network is to identify and quantify relationships between variables, very often in these cases through evaluation of correlations or conditional dependence on each pairwise combination of variables. Taxa abundance is determined by the magnitude of sequencing read counts, which is affected by sequencing depth and varies from sample to sample. Typically a sum constraint is imposed (1 for probability, 100 for percentage or 10^6^ for part per million). Thus, data are described as proportions and the sample depth is irrelevant, referred to as compositional data [5]. The only reliable information that can be extracted from these NGS data are the ratio between the parts [6] and, as noted by Pearson at the end of 19th century [7], proportional data can generate spurious correlations between measurements that are in reality uncorrelated. From a purely formal point of view, the data lie on a simplex and it can be extremely dangerous to use Euclidean metrics for proximity and correlation estimations. These compositional biases can be devastating in some datasets, and the heterogeneity (referred to as E, see Materials and Methods) together with the dimensionality D can be good predictors of their strength. The more counts concentrated in a few taxa, the greater the biases and conversely, when counts are more homogeneously distributed over samples the biases decrease. In the extreme case of only two variables, the correlation between the relative abundances will always be −1, expressing the worst possible scenario. Furthermore, a large part of taxa in the NGS experiments are under the detection limits of the sequencing techniques producing really sparse abundances matrices. It’s really common to find datasets where more than 70 − 80% of species are undetected and typically it is assigned the value of 0 and this introduces further bias in the correlation measures. The undetected species are not to be interpreted as the absence of that species but rather as a missing value in which we have no further information.

Recent methods such as Sparse Correlations for Compositional data (SparCC) [8], Sparse and Compositionally Robust Inference of Microbial Ecological Networks (SPIEC-EASI) [9] or Proportionality for Compositional data (Rho) [10] have been developed and all make extensive use of the compositional theory introduced by Aitchison [6]. He provided a family of transformations to treat this kind of data, called log-ratio transformations. For each sample, the counts are expressed with respect to a reference in order to compare them and then the logarithm is applied. Almost always the choice is the centered log-ratio transformation (CLR) where each element is divided by the geometric mean of the sample in a logarithmic scale. Thanks to this operation, the data are mapped on the Cartesian plane even if there are still limitations: the sum of all the values on each sample is always equal to zero and is equivalent to saying that the points are restricted to a D − 1 dimensional hyperplane, introducing dependencies between data. In the work mentioned above, the study focused on characterizing performance of the measurement on real and synthetic data. We focus on quantification of biases introduced on the correlations due to the CLR transformation or the sum constraint (L1) on each sample. To characterize these biases we generated synthetic data based on the ‘Normal to Anything’ approach: this method allows generation of random variables with arbitrary marginal distributions from multivariate normal variables with desired correlation structure.

## Materials and methods

### HMP2 16S Human Gut Data

Human Microbiome Project v2 (HMP2) OTU counts and their taxonomic classification were obtained from a project studying the microbiomes of healthy and prediabetic subjects over a period of up to four years [11]. The entire dataset is composed of 1122 samples and 1953 OTUs collected from 96 different subjects associated with the healthy information in each sample. To obtain a more homogeneous dataset for the following analyses we chose a subset of 51 samples belonging to a single healthy subject (69-001) with the highest number of samples.

### Network Reconstruction Methods

Many methods were developed in recent years to estimate the relationship structure of metagenomics data. We choose to focus on four among the most recent and commonly used approaches in this field. First, SparCC [8] estimates Pearson’s correlation assuming data have a sparse correlation structure. Rho [10] is based on the proportionality concept to avoid possible spurious correlations between compositional data. Finally, SPIEC-EASI [9] is a graphical model inference framework which again assumes that the underlying ecological associations are sparse and offers two different criteria to reconstruct the network i) sparse neighborhood selection MB [12] and ii) inverse covariance GLASSO [13]. In addition, we use Pearson’s correlation after data transformation (Pearson+CLR) to be compared with these more sophisticated methods.

### *α*-Heterogeneity

The *α*-heterogeneity *E* of a dataset is defined as the mean value of the α-diversity, calculated by the Shannon index, over all the samples normalized with respect to the dimensionality *D* (i.e. number of taxa). Given a dataset **X** ∈ *𝒩*^*N,D*^ composed of N distinct samples 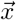 of dimension *D*, we adopt the normalized Shannon index as:

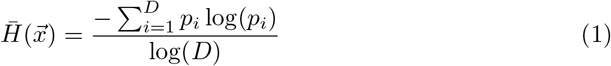

with *p*_*i*_ corresponding to the i-th taxa relative abundance in the sample, and with the *α*-heterogeneity of the dataset calculated as:

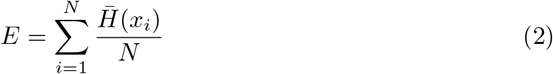

### Synthetic Data Generation

To produce artificial datasets with required characteristics (dimensionality *D*, correlation structure defined by a matrix *R*, heterogeneity *E*) we developed a toy model which makes use of the Normal to Anything (NorTA) approach to generate an arbitrary multivariate distribution with the desired correlation structure *R* as input relying on the copula functions theory [14]. A matrix *U*_*N*_ with dimensions *N*× *D*, with *N* the number of samples and *D* the dimensionality is generated from a multivariate normal distribution with zero means and *D* × *D* correlation matrix *R*. On every marginal distribution 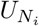 the normal cumulative density function (CDF) is applied. The inverse of cumulative function of any distribution is calculated to obtain the data used in the study. The aim is to observe correlation bias due to L1 and CLR normalizations for changes in sample dimensionality and heterogeneity. The choice of dimensionality is inherent in the NorTA method and it depends on the dimension of the input multivariate normal distribution. To tune the sample heterogeneity, we operated by tuning the magnitude of a single variable with a multiplicative factor, since very often in real data the samples are extremely heterogeneous with most of the counts contained in a few species. In summary, the model works as follows:

1. Generation of the Gaussian multivariate distribution *U*_*N*_ with dimension *D* and correlation matrix *R* as input.
2. Transformation of the data into the desired distribution *U*_NorTA_ and the calculation of the new correlation structure *R*_NortTA_. It should be noted that the correlation structure can be totally modified when strongly non-Gaussian distributions are chosen.
3. Tuning the heterogeneity of the resulting dataset applying a specific multiplicative factor to a single variable.
4. Normalization and calculation of the correlation matrix for the two normalizations L1 and CLR.

### Hurdle Truncated Log-Normal Distribution

A crucial aspect in the generation of plausible synthetic data is the choice of the marginal distribution that best describes the trend of the real data. We adopt an hurdle model that splits into different processes the zero and non-zero values [15]. It is assumed that the distribution of non-zero values in logarithmic scale follows a Gaussian with values left-censored in zero (htrlnorm):

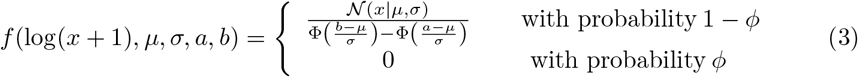

where *ϕ* is the percentage of zeros, *μ* and *σ* the mean and the standard deviation, *a* and *b* are the left and right interval limits always fixed for *a* = 0, *𝒩* (*x* | *μ, σ*) is the Gaussian probability density function and Φ its cumulative distribution. Generally the negative binomial zero inflated distribution (zinegbin) is used to model the abundances [16] even if it does not replicate very well the overdispersion and skewness of real distributions [17]. We show the comparison between them in the supplementary section S1 Appendix.

## Results

### Methods Comparison

We compared the results obtained with SPIEC-EASI (both the GLASSO and MB criteria), SparCC and Rho with respect to the Pearson’s correlation on CLR transformed data. For this purpose, we selected 51 samples of subject 69-001 in healthy condition from HMP2. We filtered the data in order to remove the rarest OTUs keeping those with a prevalence greater than 33% and the taxa with the median of the non-zero values greater or equal to 5. The comparison between the Pearson+CLR with SparCC and Rho is direct, since these three methods produce values between -1 and 1: the scatter plot of the respective correlation values is ≈ 0.99 for both, in very good accordance to a *y = x* linear relationship (Fig. 3). Since SPIEC-EASI produces a binary output in terms of conditional independence between each pair of variables, we consider the histogram of Pearson+CLR values and overlap bins corresponding to couples of variables significantly associated by SPIEC-EASI imposing a threshold on overall stability equal to 5%. Most of the significant links for SPIEC-EASI are associated to high absolute values of Pearson+CLR, with a more marked similarity for GLASSO criteria (Fig. 3). We also compared the resulting networks of the two SPIEC-EASI methods with the one obtained using the Pearson+CLR method. We get the significant links making a threshold on the p-value associated to the Pearson correlations adjusted with the Bonferroni multiple testing set to 0.05. We can observe the similarities between networks as quantified by the links shared between them (Fig. 4). The Pearson+CLR and the GLASSO criteria return almost identical results, with more visible differences for the MB method. Finally, we observe that all the obtained networks are more skewed towards positive links (Fig. 4).

**Fig 1.**
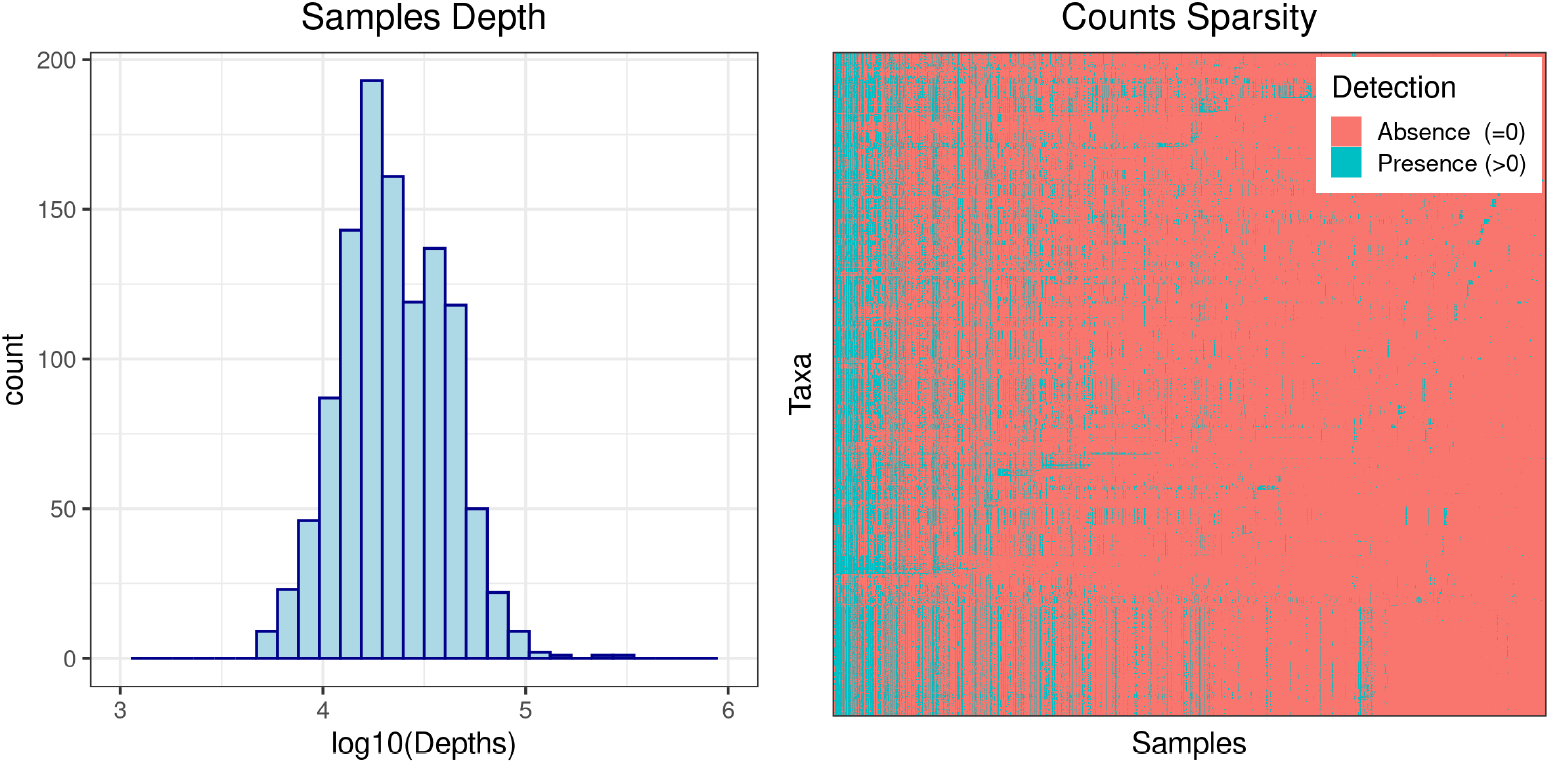
NGS Data Features. Typical characteristics of NGS data as observed in the HMP2 dataset studied in this paper. On the left the histogram in log_10_ scale of sample depths which can range from 10^4^ ≈to ≈ 10^6^. On the right the image represents the taxa detected in this dataset, over 87% of whose counts are null, equivalent to not detected taxa.

**Fig 2.**
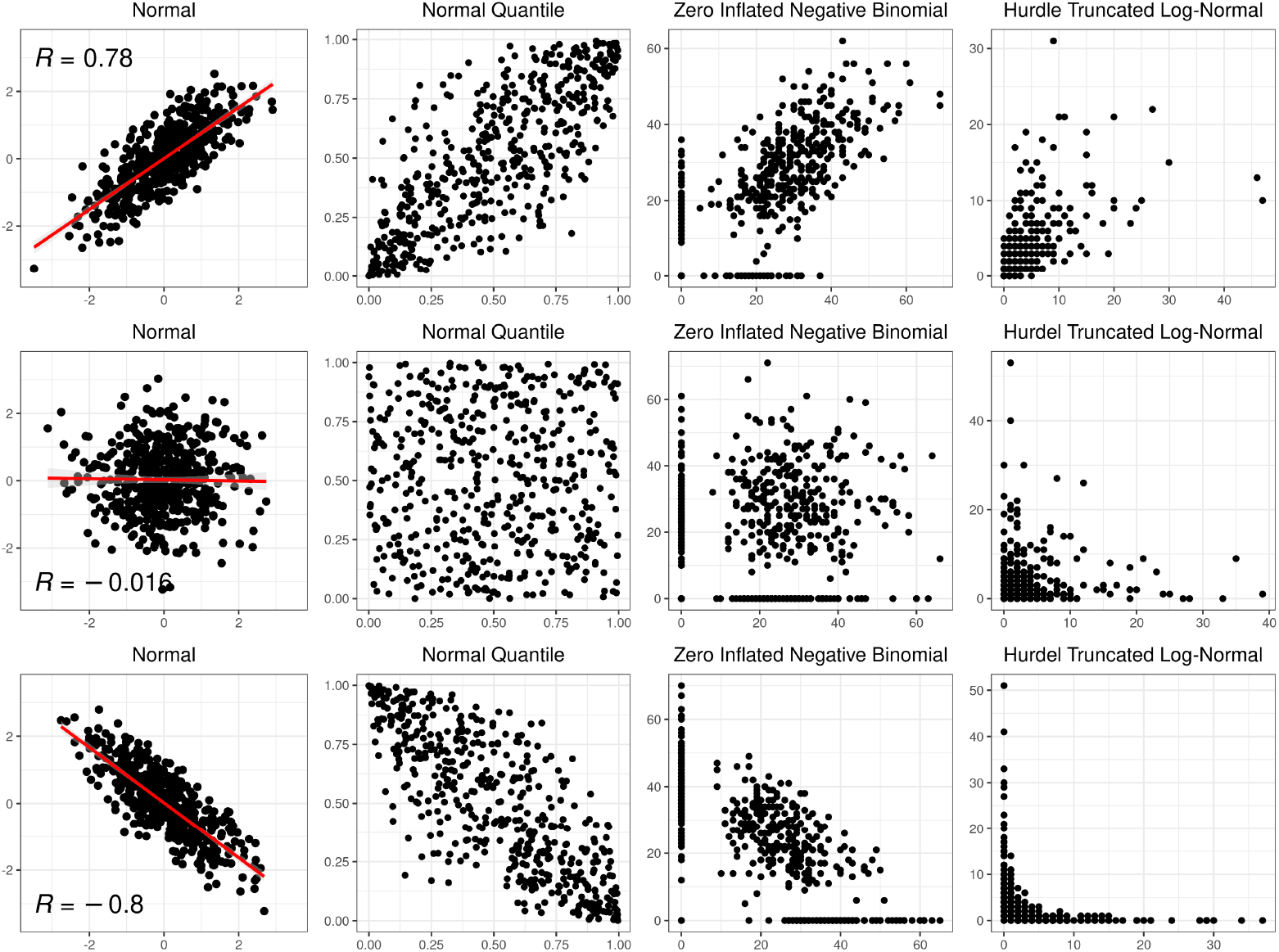
Bivariate Illustration of the “Normal to Anything” Approach. Example of Normal to Anything method on a simply bivariate case. First column: normal generate variables with different values of correlation ≈− 0.8, 0, and 0.8. Second column: non parametric remapping over quantiles. Third and fourth columns: marginal distributions for zero inflated negative binomial and hurdle truncated log-normal distributions, respectively.

**Fig 3.**
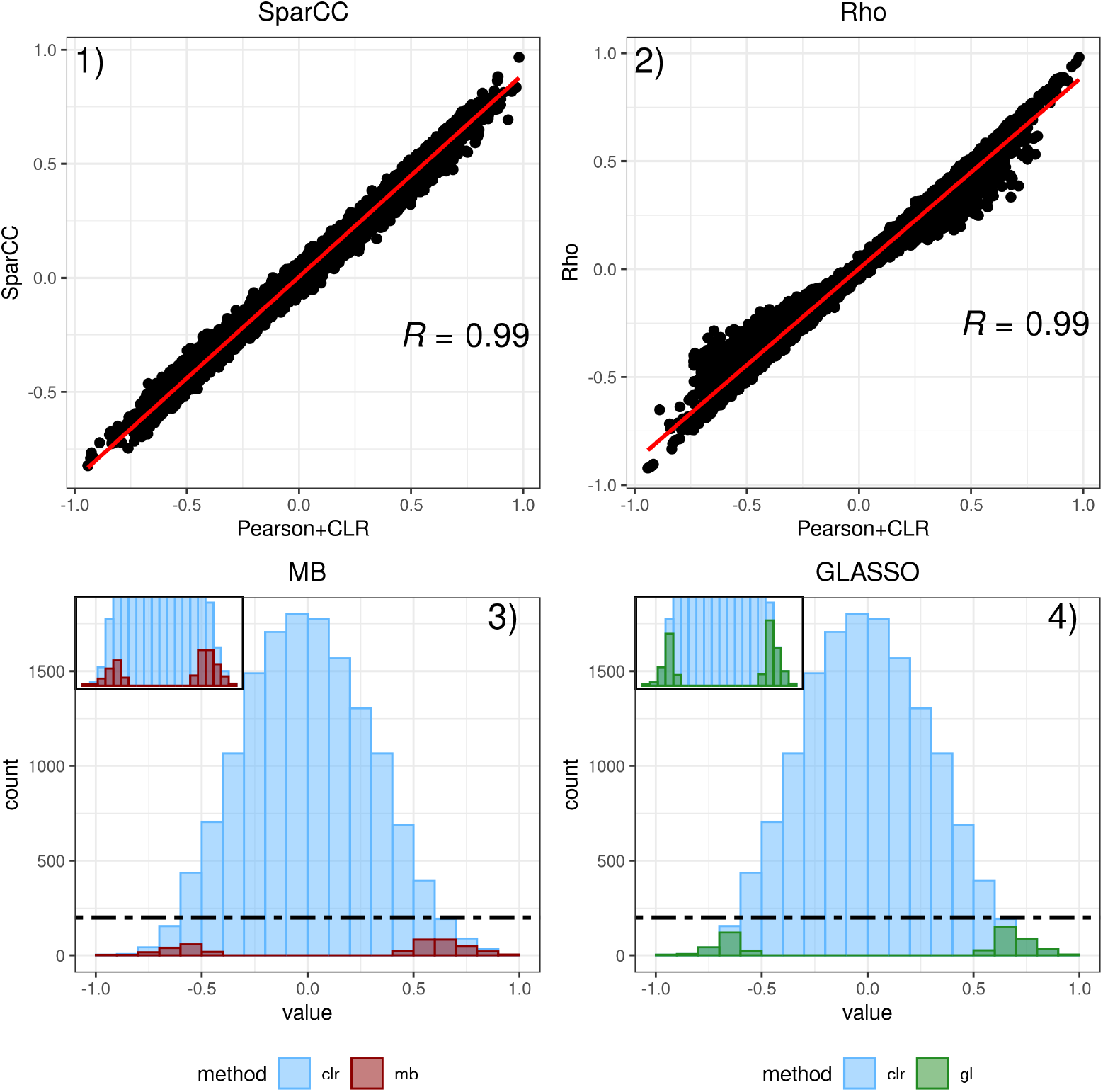
Methods Comparison. Comparison of SparCC, Rho, SPIEC-EASI MB and SPIEC-EASI GLASSO with respect to the Pearson Correlation on CLR transformed data. The scatterplot in subfigures 1 and 2 clearly show the high similarity of the two methods with Pearson correlation. Subfigures 3 and 4 are histograms of the Pearson correlation coefficients (blue bins) overlapped with the SPIEC-EASI significant links (red for MB and green for GLASSO).

**Fig 4.**
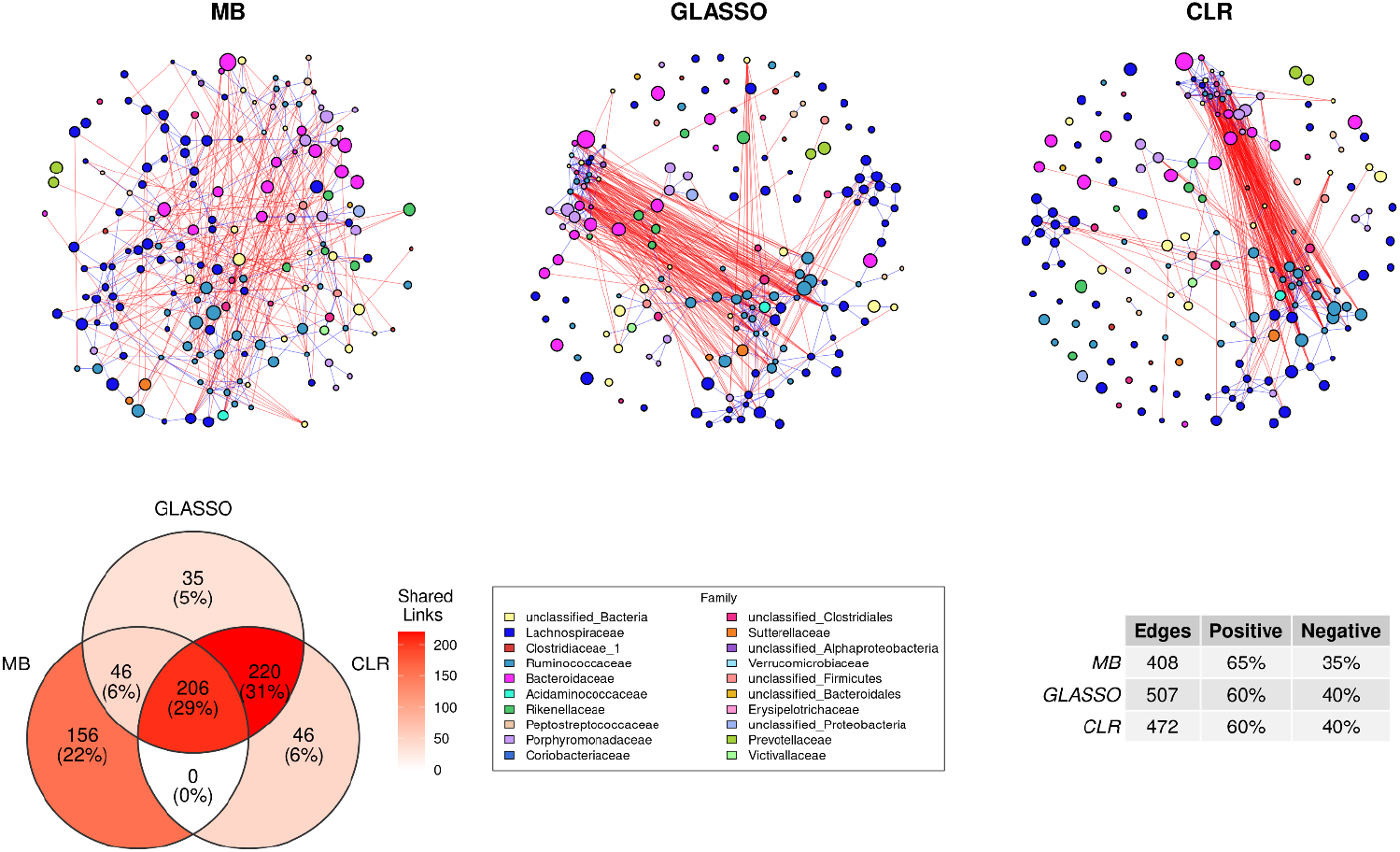
Network Comparison. Top: networks obtained with SPIEC-EASI MB, SPIEC-EASI GLASSO and Pearson+CLR methods, respectively. Vertices represent taxa, with sizes proportional to the mean abundance and colors associated to the family they belong to. Blue and red links: significant positive and negative correlations, respectively. Bottom: Venn diagram of shared links (left); legend of families represented in the above networks (centre); number of links and percentage of positive and negative links (right).

### Transformation Effects

To understand and quantify the biases caused by L1 and CLR transformations, we generate data with varying α-heterogeneity *E* and dimensionality *D*. We quantify the difference between the correlation matrix *R* generated from normally distributed data and the resulting correlation matrices *R*_*L*1_ and *R*_*CLR*_ as follows:

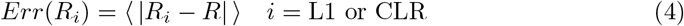

averaged over all matrix values (*Err* ∈ [0, 2]). The behaviour of the two normalization procedures is significantly different (Fig 5): L1 data are mainly affected by dataset heterogeneity, while CLR data are independent of dataset heterogeneity and the distortions decrease rapidly with dimensionality *D*. We can thus state that as the number of taxa increases, the compositional effects decrease becoming negligible for *D* > 80.

**Fig 5.**
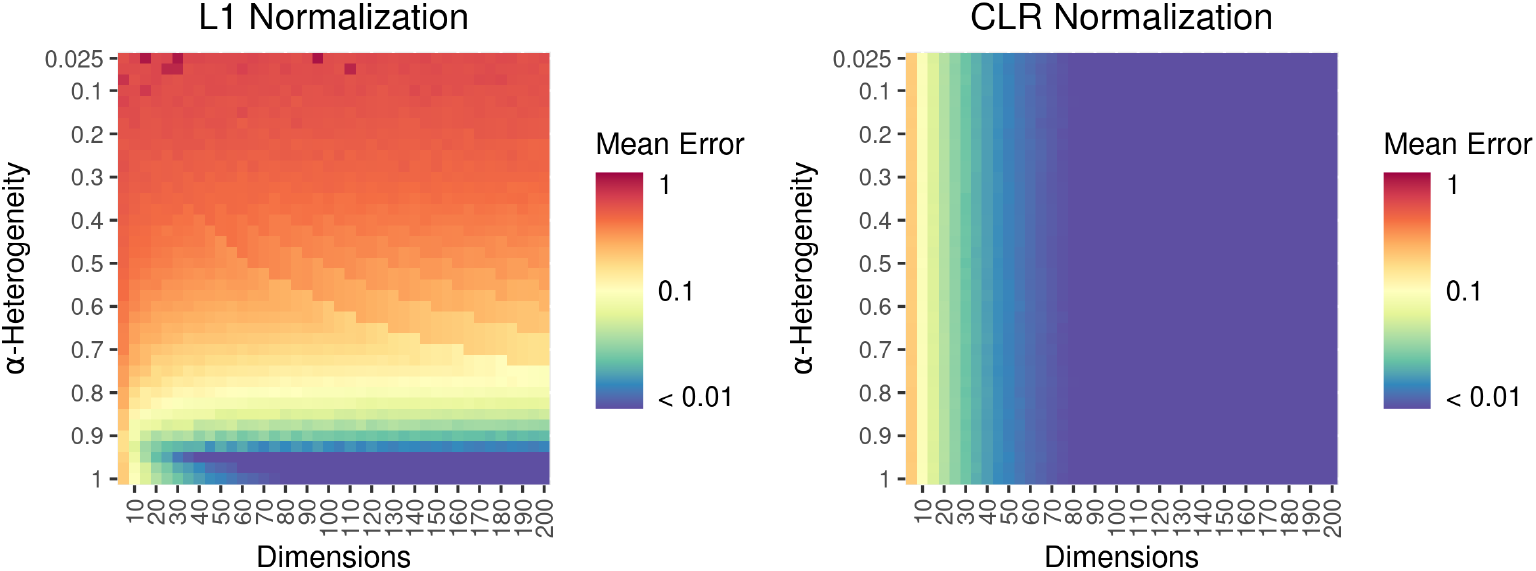
Heatmap of *Err* as a function of α-heterogeneity and dimensionality for L1 normalization (left) and CLR normalization (right).

### Effects of data sparsity

To describe the effects of a large number of below-threshold abundance taxa (zero-value measurements) we study a simulated dataset of 100 taxa, in order to minimize normalization biases due to low dimensionality *D* as described in previous section, generated from the hurdle truncated log-normal distribution, varying the correlation strength between two taxa (both positive and negative correlations) and the percentage of below-threshold observations (Fig. 6).

**Fig 6.**
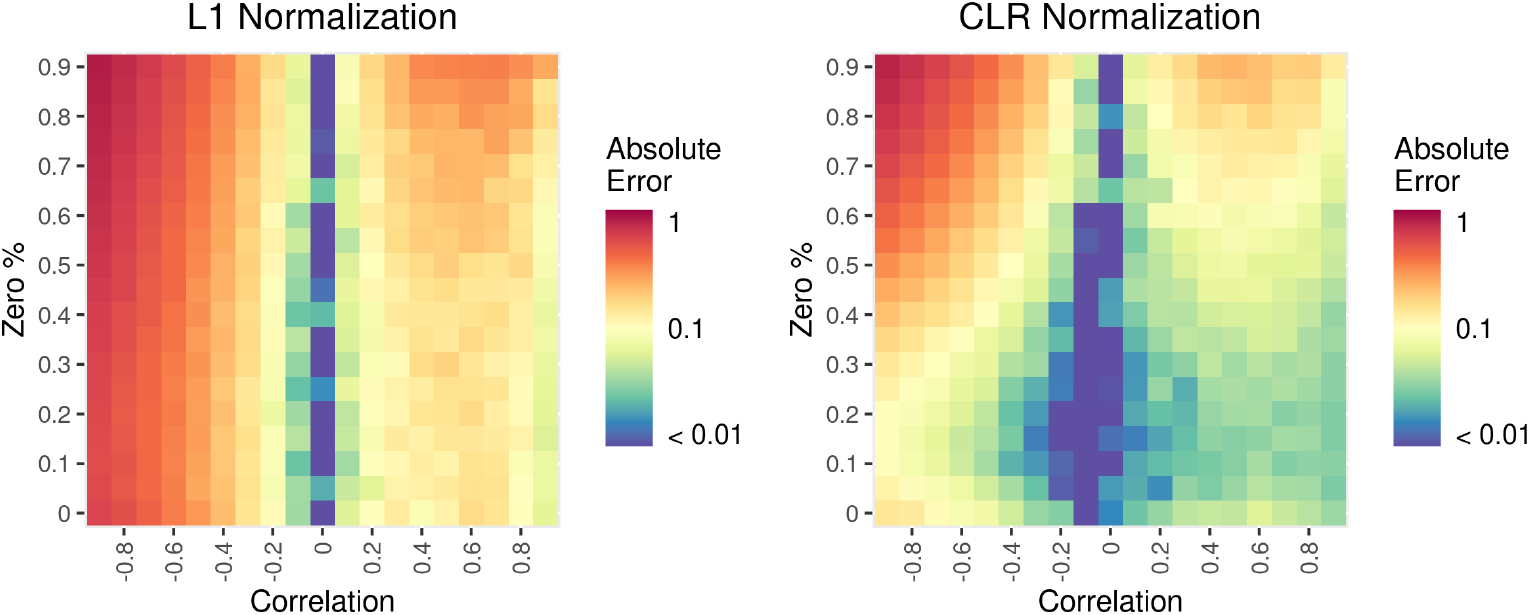
Absolute error on correlation as a function of initial correlation values before NorTA transform and of zero count percentage: left L1 normalization, right CLR normalization.

We observe a very different behaviour with respect to the sign of the correlation: the largest errors are associated to negative correlations in general, and mostly for the L1 normalization case. On the other hand, CLR normalization is more successful in retrieving negative correlations values when the amount of zero values does not exceed 30%, while for positive correlations, up to 60% of zero values is tolerable.

## Conclusion

The network analysis framework can help to gain deeper insight in metagenomic studies, for example to characterize community structure in different conditions. A very simple and common approach is to reconstruct networks based on measurement correlation, but the compositional nature of metagenomics data poses some challenges, and specific methods have been developed to overcome this issue. In this work we thoroughly explore the behaviour of different available methods, as a function of several factors, such as the dimensionality and heterogeneity of the dataset, including the data sparsity, very common features of these type of data obtained from NGS sequencing independently of the technique (eg 16S or whole-genome approaches). We observe that for relatively high-dimensional data, as is the case for NGS metagenomics, many biases are easily removed or attenuated by simply applying compositional data normalization procedures, and after this proceduree the classical Pearson’s correlation can provide results very similar to other existing algorithms specifically developed to deal with metagenomics data. Data heterogeneity (ie the disproportion of abundances within one sample) can affect correlation, but this effect is negligible for CLR normalization. The percentage of zero measures in the dataset, related to low-abundance taxa, can significantly affect correlations, producing a bigger distortion on negative correlations that can result underestimated. Since the bias introduced by zero measures cannot be completely removed, in general a tradeoff will be necessary between minimizing correlation distortions and keeping low-abundance species in the analysis.

## Supporting information

Data and codes are available at https://github.com/Fuschi/Correlation-Biases-on-Metagenomics-Data. We also developed a lightweight R package to generate synthetic metagenomic data as explained in the Materials and Methods section available at https://github.com/Fuschi/ToyModel.

### S1 Appendix

#### Hurdle Truncated Log-Normal Distribution Benchmark

Comparison between zero-inflated negative binonial and hurdle truncated log-normal distribution to model real NGS data from HMP2 dataset.

## Acknowledgments

We are grateful for discussions with Jethro Johnson, Farzaneh Rastegari, and Juliana Alcoforado Diniz during preparation of this manuscript.

